# A Comprehensive Resource for Exploring Antiphage Defense: DefenseFinder Webservice, Wiki and Databases

**DOI:** 10.1101/2024.01.25.577194

**Authors:** F. Tesson, R. Planel, A. Egorov, H. Georjon, H. Vaysset, B. Brancotte, B. Néron, E. Mordret, G. Atkinson, A Bernheim, J. Cury

## Abstract

In recent years, a vast number of novel antiphage defense mechanisms were uncovered. To facilitate the exploration of mechanistic, ecological, and evolutionary aspects related to antiphage defense systems, we released DefenseFinder in 2021 (Tesson et al., 2022). DefenseFinder is a bioinformatic program designed for the systematic identification of known antiphage defense mechanisms. The initial release of DefenseFinder v1.0.0 included 60 systems. Over the past three years, the number of antiphage systems incorporated into DefenseFinder has grown to 152. The increasing number of known systems makes it a challenge to enter the field and makes the interpretation of detections of antiphage systems difficult. Moreover, the rapid development of sequence-based predictions of structures offers novel possibilities of analysis and should be easily available. To overcome these challenges, we present a hub of resources on defense systems, including: 1) an updated version of DefenseFinder with a web-service search function, 2) a community-curated repository of knowledge on the systems, and 3) precomputed databases, which include annotations done on RefSeq genomes and structure predictions generated by AlphaFold. These pages can be freely accessed for users as a starting point on their journey to better understand a given system. We anticipate that these resources will foster the use of bioinformatics in the study of antiphage systems and will serve the community of researchers who study antiphage systems. This resource is available at: https://defensefinder.mdmlab.fr.

## Introduction

In the last few years, a considerable number of newly discovered antiphage defense mechanisms have come to light ^1^. The most widespread mechanisms, Restriction-Modification and CRISPR-Cas systems, target foreign nucleic acids^2,3^. However, recent discoveries have revealed an important diversity of molecular modalities by which bacteria defend themselves against phages. This diversity of mechanisms includes nucleotide depletion 4–10, membrane disruption11–14, and production of antiviral molecules^15^. Importantly, many defense mechanisms remain unknown.

The discovery of diverse antiphage systems not only provides new insights into bacterial immunity mechanisms but also transforms the exploration of interactions between phages and bacteria and microbial evolution. Evaluating the impact of different defense systems in naturally occurring *Vibrio* isolates has revealed that a rapid turnover of a few mobile genetic elements encoding defense systems can completely alter their susceptibility to phages^16,17^. Ongoing research is delving into the broader phylogenetic scale to explore the role of defense systems and their potential implications for phage therapy strategies^18,19^. Furthermore, examples indicate that bacterial defense systems can be co-regulated and operate synergistically in multi-layered defense strategies^20^. Beyond experimental approaches, there is a growing interest in identifying the antiphage systems encoded in diverse species or environments to understand the diversity of antiviral strategies employed by bacteria.

To investigate mechanistic, ecological, and evolutionary questions related to antiphage defense systems, we created in 2021 DefenseFinder^21^, a tool to detect known antiphage defense systems systematically. At the time of publication of DefenseFinder v1.0.0, the tool encompassed 60 systems. In the last 3 years, the number of antiviral systems included in DefenseFinder has grown to 152. The increasing number of systems has been accompanied by emerging challenges in comprehending DefenseFinder results biologically. For non-specialist users, the analysis can be quite intricate, demanding a high level of knowledge in this rapidly evolving field.

To bridge this gap, we decided to create a website dedicated to defense systems, encompassing an improved version of a web service to run DefenseFinder and diverse databases to illuminate and increase the understandability of bioinformatic detection of antiphage defense systems. Here, we provide the release of 92 new defense systems models and a software update as well as 3 databases. 1/ A collaborative knowledge base (wiki) summarizing information on known defense systems. 2/ A structure database, with experimentally determined and AlphaFold2 predicted structures. 3/ A precomputed database of DefenseFinder results in over 20,000 complete genomes. Besides, we designed the website to keep it as up-to-date as possible. Wiki pages are easily editable by anyone and reviewed by experts in the field before publication on the website. They can also be edited automatically to generate sections that aggregate data, such as the phylogenetic distribution of a system. All those website components are also integrated within the DefenseFinder webserver output to easily find information on a system found in a genome.

## Results

### Updates of DefenseFinder program

To improve detections performed by DefenseFinder, we updated both the program and the models. Since the release of DefenseFinder models v1.0.0, we have constantly updated the defense system models included in DefenseFinder by adding a total of 92 defense systems and 132 subsystems (**Figure 1A**). Among those systems, 63 were discovered and described after releasing the first version of DefenseFinder models. Other systems were missed in the first version and are now added after a deeper literature review of the field. Those systems encompass the “Abi” group discovered between 1990 and 2006. DefenseFinder models v1.0.0 included 3 of these systems, while the latest version v1.2.3 detects 22 Abi systems^22^. Beyond Abi systems, we also added SanaTA ^23^, MazEF ^24^, antiphage defense systems identified in mycobacterium prophages ^25,26^ and pAgo ^27^. The defense system named “Rst_DprA-PPRT ’’ ^28^ was renamed “ShosTA” as it was initially discovered in 2013 ^23^ but without phage activity testing. Finally, for Lamassu-Fam^29^, we separated the systems into different subsystems according to the evolving literature on the topic. Models were also adapted to the latest version of MacSyFinder^30^.

**Figure 1.**
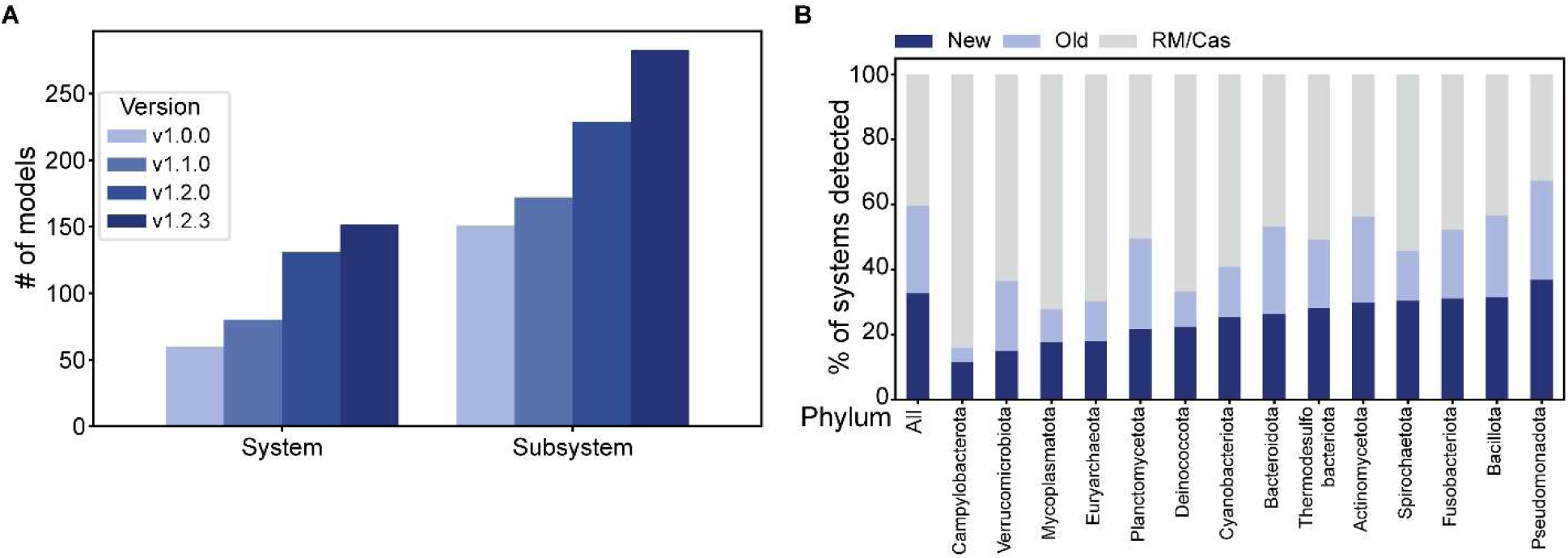
Evolution of DefenseFinder models. **A.** Evolution of the number of defense systems across the different versions. **B.** Proportion of the currently detected systems which were added in v1.0.0 (Old and RM/Cas) or in more recent versions (New) of DefenseFinder models across different phyla of prokaryotes.

To evaluate the proportion of newly detected systems, we ran DefenseFinder v1.2.3 on the RefSeq complete genome database of prokaryotes (Bacteria N = 22,422; Archaea N = 381). We found a total of 152,386 different systems, among which 33% correspond to systems present in v1.2.3 but absent from v1.0.0 (**Figure 1B**). Without restriction-modification and CRISPR-Cas systems, which are by far the most abundant systems, newly added systems now represent 54% of the detected systems (41-73% across phyla) (**Figure 1B**).

The previous version of DefenseFinder used a “profile coverage” parameter of 0.1. This value was chosen to detect defense systems when the protein was split. This low coverage threshold was balanced with a high score threshold for single-gene systems and the necessity to find the different components of the system colocalizing in the genome for multi-gene systems. We changed that threshold to 0.4 as a default value in the latest version. This threshold was set to maintain the possibility of finding incomplete genes or a gene with a domain replacement while removing very low coverage off-target hits. Users can still manually change the coverage threshold (--coverage-profile) to decide on how conservative detection should be depending on the biological problem at hand. For ease of use, we also added the possibility to directly input a nucleic acid fasta file. We use pyrodigal v3.0.1^31^ to identify and translate the coding regions, which are then processed by DefenseFinder. Finally, we added non-regressive tests in a continuous integration pipeline to make the development of future features more robust against introduction of bugs.

### An improved web service

DefenseFinder is a Python program running in the command line. To enhance the software’s usability, DefenseFinder has been available as a web service since its launch. Some useful features were missing, such as the possibility of accessing previous analyses. We updated the web service to add new features: a simple interface for depositing fasta nucleic acid or fasta amino acid entries on the "Home" page (**Figure 2A**). Results of previous analyses are now displayed in the “Analysis” page. On this page, jobs can be accessed and renamed at convenience for 1 month without any action on the web service. All the different DefenseFinder outputs are displayed in dynamic tables that can be easily downloaded. The results are displayed in a genome browser to better visualize the hits in their context (**Figure 2B**). All results are displayed, including orphan HMMs, which do not form a system. Those orphan HMMs should be used cautiously and analyzed using their score and profile coverage. From the result table, getting more information on a given system is now easier via a link to the collaborative knowledge base. Importantly for the community, the latest version of DefenseFinder has been packaged in Galaxy ^32^ and can be run on any Galaxy instance.

**Figure 2.**
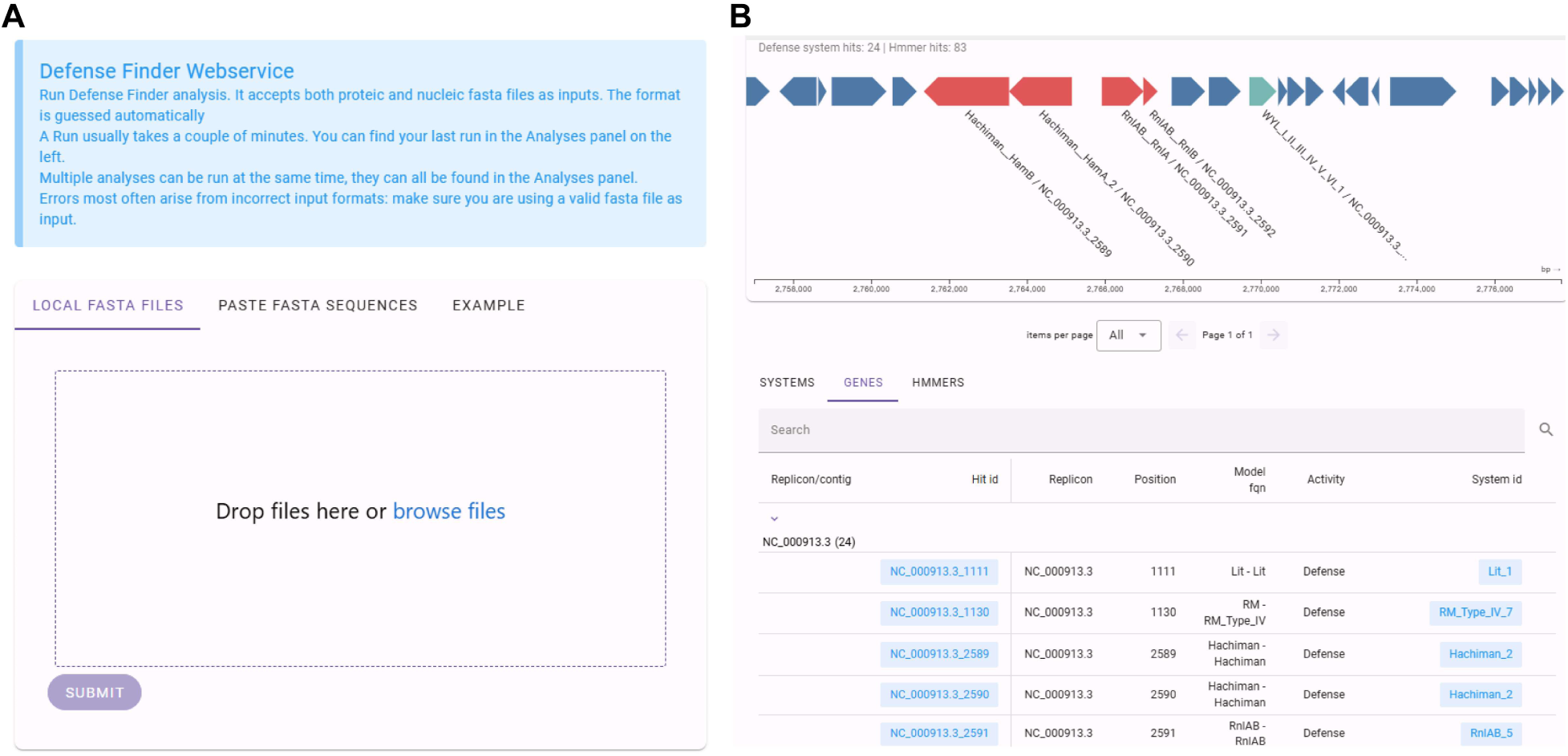
Architecture of DefenseFinder Webservice. **A.** Homepage of the DefenseFinder webservice. **B.** Results page of the webservice. Three different output tables and a visualization of the chromosome with the different defense system hits are available.

### A collaborative knowledge base for Defense systems

DefenseFinder output constitutes 3 tables (defense_finder_systems, defense_finder_genes, and defense_finder_hmmer) containing respectively one line per detected system, detected genes inside a system, and defensive HMM profiles hits. Given the large (and ever-growing) number of defense systems, this output can be hard to analyze. Thus, we decided to create a participative knowledge base of defense systems available at https://defensefinder.mdmlab.fr/wiki/. This website aims to provide information to understand better bioinformatic predictions provided by DefenseFinder and to allow a dynamic, up-to-date sharing of knowledge on antiphage systems. The wiki provides a few pages on general concepts of the field and a page per defense system detected by DefenseFinder. The different pages are accessible directly from the results of the DefenseFinder web service or can be explored individually and are all summarized in a table containing the main references, part of their mechanism (sensor, activator, effector) when this information is available in the literature and the Pfam domain present in the system (**Figure 3A**).

**Figure 3.**
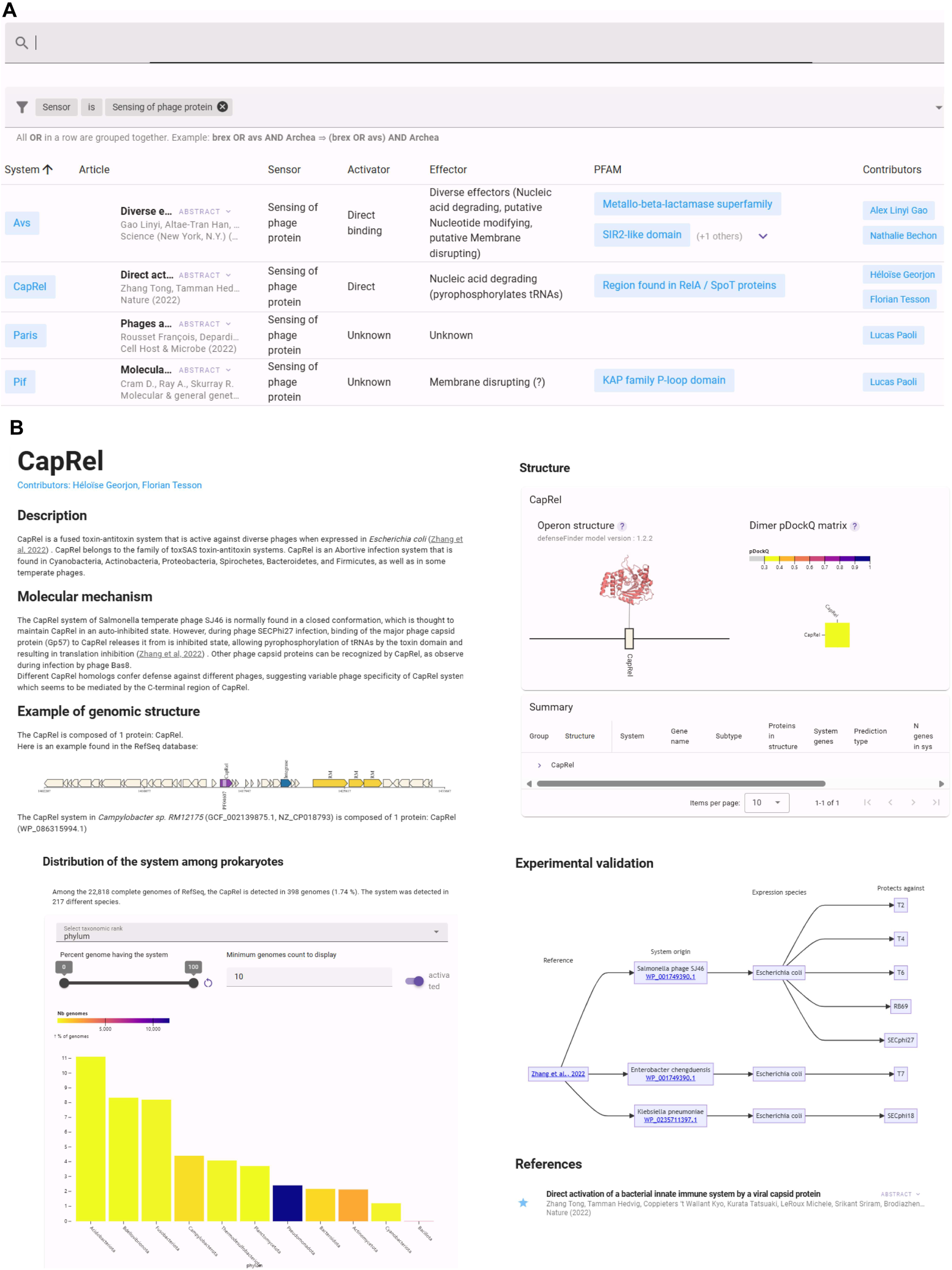
Architecture of the DefenseFinderWiki collaborative knowledge base. **A.** Table presenting all the different defense systems. The table can be filtered by system, type of sensor/activator or effector and Pfam inside the system. **B.** An example of one page of the knowledge base covering the CapRel defense system.

Each defense system page is organized into distinct sections (**Figure 3B**). 1/ **Description**: summarizing simple information such as the system’s discovery, protein names, and known domains. 2/ **The molecular mechanism**: summarizing known mechanisms and highlighting instances where the mechanism remains unknown. 3/ **Genomic architecture**: a detailed breakdown of system components and associated domains is provided, accompanied by examples of genomic loci organization. 4/ **Distribution among prokaryotes**: the system distribution across phylogenetic phyla using DefenseFinder results on the RefSeq complete genome database. 5/ **3D Structure of the system**: showcases both experimentally validated 3D structures (if available) and predicted structures using AlphaFold2 ^33^ for validated systems. Foldseek ^34^ results starting from the given predicted structure are also precomputed and available. 6/ **Experimental validation**: shows the tested system, the expression organism and against which phage(s) it was shown to be effective. 7/ **References**: relevant publications (denoted with a star) are included to facilitate a deeper understanding of each system.

Each system featured in DefenseFinder models v1.2.3 has a dedicated page on the knowledge base. These pages were populated through collaborative efforts, including a hackathon organized internally and contributions from other researchers from the community. Recognizing the rapid pace of defense system discovery and mechanism characterization, we opted for a collaborative approach by making the website open for updates. To foster a collaborative framework, we based the website on a Gitlab repository, with pages written in Markdown and designed to be easy to use by people that are not Git experts. To modify and add information to wiki pages, contributors can click on “Edit a page” (found at the bottom of every page), add some content (and add themselves as contributors) and create an automatic merge request that will be verified by at least one expert in the field before being merged. When describing a new system, the discoverers are encouraged to request the creation of a new page to describe it with all the necessary information.

### Precomputed results of DefenseFinder on 22,738 complete genomes

Running DefenseFinder on a large database is time and resource-consuming; thus, we created a precomputed database. DefenseFinder v1.1.0 with models v1.2.3 was run on the complete genomes database from RefSeq (From July 2022, Bacteria N = 22,422, Archaea N = 381). A total of 152,386 defense systems were detected from 152 different defense systems (264 subtypes). Those precomputed results can be visualized on the “RefSeq DB” tab of the DefenseFinder website: https://defensefinder.mdmlab.fr/wiki/refseq/. Results are displayed in an interactive table, allowing research by system, accession, or taxonomy (**Figure 4A**). This table is linked to interactive graphs displaying the relative abundances of the system in the phylogeny and distribution of systems (**Figure 4B and 4C**). Results (tables and graphs) can be downloaded from the website either as a whole or only a subset using different filters. Sequences of detected proteins are accessible through a link to their NCBI protein page.

**Figure 4.**
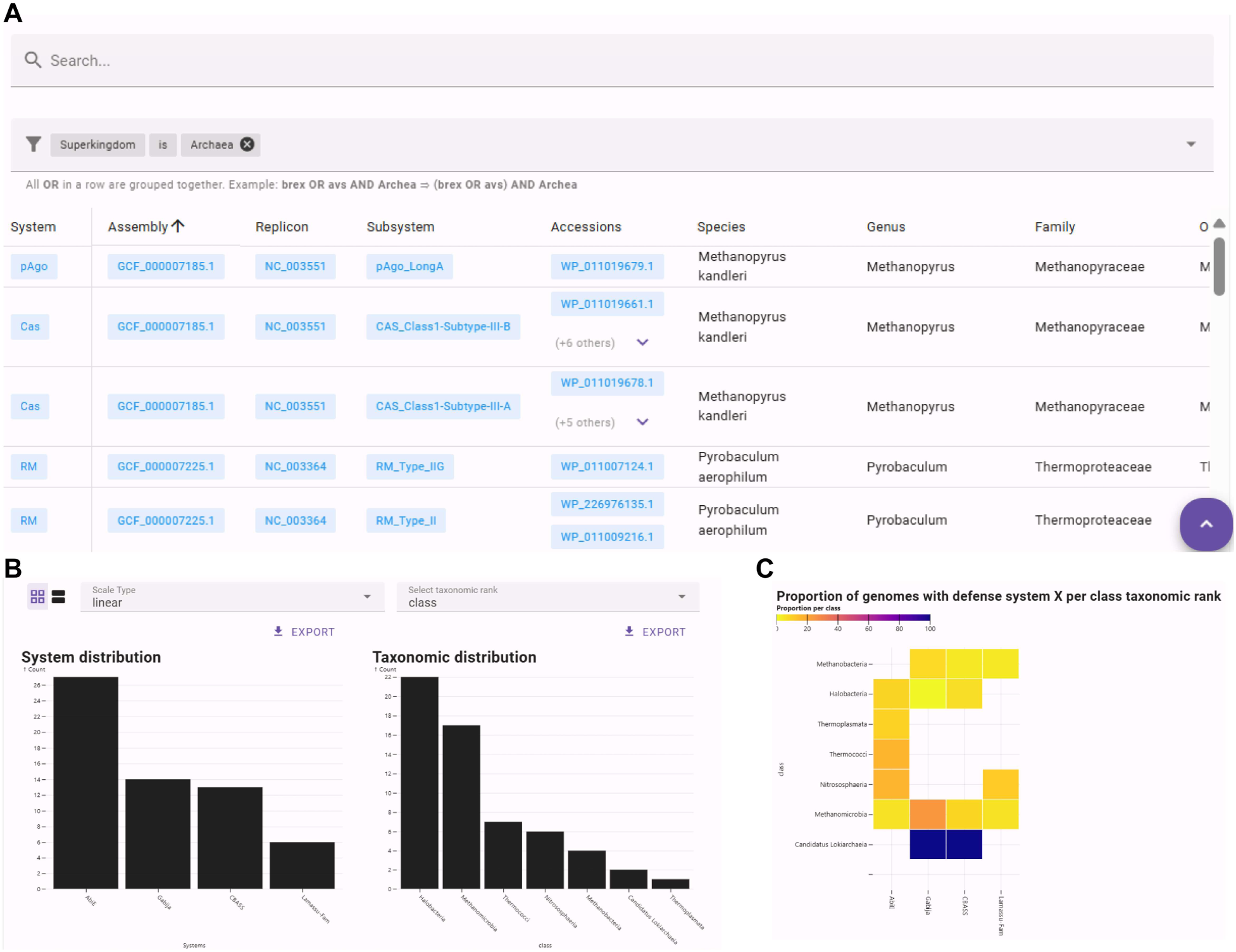
DefenseFinder results database. **A.** Main table used to search through the DefenseFinder results. Searching through this table can be done using only keywords or by using a combination of conditions on the system, the phylogeny, or the name of the replicon. **B.** Interactive bar charts of the count of each system on the left and the count of each taxonomic order reactive to the filter in the main table. **C.** Interactive heatmap displaying the count of each system in each taxonomic order reactive to the main table. The taxonomic level can be modified from superkingdom to the species level for both the barchart and heatmap.

### Precomputed 3D structure predictions of defense systems

Many mechanisms of the newly described systems remain to be elucidated. Multiple studies were conducted to elucidate such mechanisms^5,9,13,35–38^. Often, the 3D structure of the system was an important step towards understanding its mechanism^4,8,10,39–41^. Recent developments in structure predictions, such as the development of AlphaFold2 or ESMFold^33,42^, allow good-quality predictions of proteins and protein complexes. We thus created a database of predicted structures for known defense systems.

Experimentally validated proteins were retrieved for each system, and the 3D predicted structure was computed using AlphaFold2. For some systems, we could not find the original protein sequence or accession of the experimentally validated system. Some subsystems (CBASS^43^, Retron^44,45^, Lamassu^29^, etc.) were not experimentally validated and are included in DefenseFinder. In such cases, we selected a different representative from DefenseFinder for structural prediction (See Methods). Recent studies demonstrated that many systems function as complexes ^4,8,10,39,40^. To provide insight on the possible complexes, all possible dimers (homo and heterodimers) were computed for each system. For each predicted complex, pDockQ scores ^46^ were computed to assess the probability of the protein-protein interaction. For several systems with three or more proteins, complexes with up to four proteins were also computed in 1:1 stoichiometry. To discover similar folds in structural databases, we ran FoldSeek^34^ using the predicted 3D structure of the monomers against the PDB ^47^ and AlphaFold Uniprot^48,49^ databases, and provided the precomputed results in the structure table.

In total, we provide more than 1,500 predictions of monomers, homodimers, or heterodimers. Results are available on the DefenseFinder structure database at: https://defensefinder.mdmlab.fr/wiki/structure/ as a table with all predicted structures present in the database (**Figure 5A**). Users can search for specific proteins or systems as well as filter all the structure by monomers, dimers, or quality statistics (pLDDT, iptm+ptm or pDockQ for multimers). Predicted structures can be visualized on the website using Mol* ^50^ with the predicted structure’s confidence score (pLDDT) (**Figure 5B**). Results can also be downloaded individually or in bulk directly on the website.

**Figure 5.**
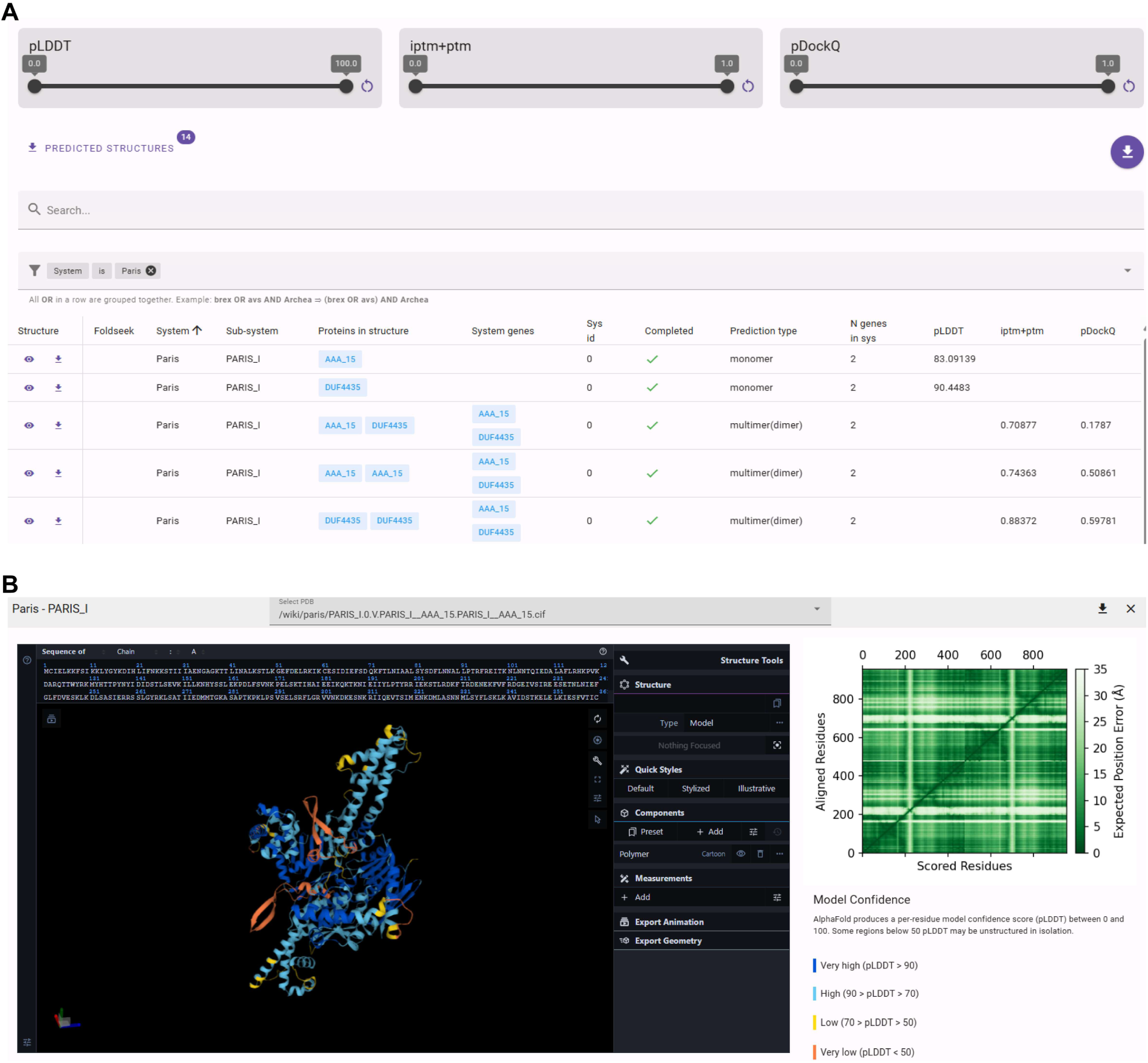
DefenseFinder predicted structures’ database: Organization of the structure database. **A.** The dynamic table to search across the 1,543 predicted structures. **B.** Example of protein structure visualization with a dimer of AriA_AAA15 from the Paris defense system.

### Conclusion and perspectives

The field of antiphage systems is growing rapidly, which can be difficult to follow for many researchers and students whose primary focus is not antiphage systems. Typically, literature reviews can become quickly outdated. At the same time, bioinformatics predictions guide many studies on antiphage systems. We offer a comprehensive resource for studying antiphage systems to lower the barrier of bioinformatics predictions and to have flexible and fast-evolving access to knowledge on antiphage systems.

The integration of DefenseFinderWiki enables the outputs produced by DefenseFinder to be linked to wiki-style pages, consolidating information on various defense systems thereby improving the understanding of the defense arsenal. Identifying defense systems can pose computational challenges, particularly when applied to large datasets. Here, we ran DefenseFinder on the RefSeq database. The resulting data is showcased through a dynamic table, complemented by visualizations depicting their distribution across the prokaryotic taxonomy. The final section of the website is devoted to protein structures, showcasing visual representations of predicted structures for all identified defense systems.

The ongoing commitment to regular updates of DefenseFinder ensures the incorporation of newly discovered defense systems, thereby enhancing the platform’s utility and adaptability within the scientific community. To match the speed of discovery, we built the platform as a collaborative database. This will allow researchers from the community to add information about previously described systems and to create new pages for undescribed defense systems. This database is easy to modify via Gitlab, even for non-git users, while taking advantage of a version-controlled system.

We will continue to develop the community aspect of the knowledge base by providing tutorials and organizing workshops to encourage people to contribute to the project. New updates will be made to increase the information on the website (new predicted structure, alphafoldDB^49^, increase in the number of genomes, sequence availability). We plan to add in a future release, a new section where users can test whether a system is related to a known one or not. If the system is new, we will provide a form to add the new system both for DefenseFinder and the website.

Overall, we are providing a new hub, gathering many resources that we hope will be useful for those exploring antiphage defense systems. This website aims to support newcomers in the field, including students, seasoned researchers, and enthusiasts of defense systems, by providing a platform to initiate work or deepen their understanding and knowledge in this domain.

## Materials and Methods

### Website development

This website comprises two distinct components: the webservice and the knowledge base (wiki and databases) each functioning separately. On the frontend, both utilize Vue.js and its ecosystem, particularly Nuxt.

The Webservice: The backend is built on top of Django and Django Ninja, designed for user-friendly and intuitive API development. It establishes communication with the Pasteur Galaxy ^51^ instance to execute the DefenseFinder tool and retrieve results. Subsequently, outcomes are stored in a PostgreSQL database accessible through anonymous sessions.

The knowledge base: Devoid of a backend, this is a fully static website implemented using Nuxt and Nuxt Content. Nuxt Content, a Git-based Headless CMS, facilitates the creation and management of static pages via Markdown or JSON files stored in a GitLab repository. This architectural approach alleviates the need for database and backend maintenance, delegating these responsibilities to GitLab, which seamlessly handles authentication, permissions, modification history, and a rich web editor. Therefore, the content is easily editable and accessible. The wiki also provides an interface for searching through large datasets (RefSeq and structure predictions) using Meilisearch (self-hosted) with filters and complex queries.

The plots are generated using Observable Plot and D3.

Both websites undergo automated deployment to a Kubernetes cluster through a GitLab CI workflow. Additionally, a custom linter can be executed against the markdown content via the GitLab web interface.

### Protein selection for structure prediction

Protein accessions of experimentally validated systems were retrieved and used for structure prediction for each system and subsystem. For several subsystems, it was not possible to retrieve experimentally validated sequences for two reasons: no protein sequences or accessions in the original paper or, it’s a subsystem with no experimental validation. For those systems, one of the best system hits from DefenseFinder was randomly selected and used for the protein structure prediction. Best hits were selected based on their hit scores and profile coverage (fourth quantile of hit score for each gene of the system and more than 75% of profile coverage).

AlphaFold2 (v2.3.1) ^33^ was used to predict protein structures, using the AlphaFold-Multimer ^52^ protocol for complexes. Monomer and complex models were sorted based on pLDDT (mean of predicted Local Distance Difference Test) and iptm+ptm (weighted sum of interface and all residues template-modeling scores), respectively. The models with the highest scores were taken for subsequent analysis. Additionally, for dimer structures, pDockQ scores (predicted DockQ) ^46^ were calculated to assess interface quality; (models with acceptable quality are considered if they have DockQ ≥ 0.23). For all monomeric structures we used FoldSeek (v5) ^34^ to perform structure similarity searches against the PDB ^47^ and AlphaFold Uniprot ^48,49^ databases.

### Precomputed DefenseFinder results

For the precomputed database, we used all the RefSeq complete genomes ^53^ of both Bacteria (N = 22,422) and Archaea (N = 381) from July 2022. DefenseFinder v1.2.0 ^21^ using models v1.2.3 was run on all genomes separately using standard settings (coverage 0.4).

### Pfam annotation

We ran hmmsearch HMMER 3.3.2 ^54^ on proteins that were detected by DefenseFinder with a coverage above 75%, to make sure they are complete, against Pfam-A database v33.1 ^55^. For each protein family of a given defense system, PFAM that hits at least 50% of the members of the family are assigned to the protein family and therefore to the system.

Pfam annotation of the example of genomic locus was done on each protein of the system using hmmsearch “with --ga_cut” argument. If two PFAM hits were overlapping in a single protein sequence, only the best hit (hit_score) was kept.

### New profile modeling

New HMM profiles were built using the same method as in the first version of DefenseFinder (see methods in ^21^. For protein with a single representative available, the first profile was made either by blasting one homolog (if only one is available) on BLAST RefSeq (nr) ^56^ database (filter with 30% identity and 70% coverage). If a first detection is available in supplementary material, the first profile is done using those proteins.

### Addition of new models

Using the architecture of MacSyFinder v2 ^57^, we added new definitions and profiles inside the previous MacSyFinder models. Inconsistencies between models were then checked and reduced: overlapping systems and systems blocking the detection of the other.

## Data availability

DefenseFinder website is available at: https://defensefinder.mdmlab.fr/. This website is free and open to all users and there is no login requirement. The source code of the wiki pages is hosted on the GitLab instance of the Pasteur Institute (https://gitlab.pasteur.fr/mdm-lab/wiki) and is under GPLv3 license.

## Acknowledgement

We are grateful to all the members of MDMLab for their help in filling most of the knowledge base pages with relevant information. We thank Nathalie Béchon, Alex L. Gao, Adi Millman, François Rousset, Avigail Stokar-Avihail and Daan Swarts who accepted to write pages on their favorite subjects. We thank the IT Department of Institut Pasteur, including Thomas Menard, in particular, for providing access to the Kubernetes cluster and initial training.

